# Information Leakage in Enzyme Substrate Prediction

**DOI:** 10.64898/2026.02.26.708291

**Authors:** Vahid Atabaigi Elmi, Roman Joeres, Olga V. Kalinina

**Affiliations:** Drug Bioinformatics, Helmholtz Institute for Pharmaceutical Research, Saarbruecken, Germany; Centre for Bioinformatics, Saarland University, Saarbruecken, Germany; Medical Faculty, Saarland University, Homburg, Germany

## Abstract

Enzymes are essential catalysts in many cellular processes. Understanding their interactions with small molecules, such as regulators, cofactors, and most importantly, substrates, is crucial for understanding the biochemical processes that occur in cells. Correctly interpreting the roles of small molecules that interact with enzymes is key to elucidating enzyme function. Recently, the field of enzyme-small molecule interaction prediction has gained more interest from computational and, especially, deep-learning methods, and numerous datasets and models with remarkable performances have been published. In this work, we critically examine one of the most popular datasets and three models trained on it, identifying leaked information that may overinflate reported model performance. We show that the inspected models are susceptible to information leakage, and their performance drops to near-random when the leakage is removed.

## 1 Introduction

Enzymes are proteins that catalyze essential biological reactions, playing key roles in metabolism, energy transfer, signal transduction, and many other cellular processes [1]. They do so by binding a small chemical molecule (substrate) and transforming it into another molecule (product), often depending on or regulated by other small molecules (cofactors, inhibitors, regulators, etc.). For example, RuBisCO (Enzyme Commission, or EC, number 4.1.1.39) is involved in photosynthesis in plants and catalyzes the fixation of atmospheric carbon dioxide into glucose and other sugars [2]. Understanding interactions between enzymes and small molecules is fundamental in biology and drug design.

Computational prediction of enzyme-small molecule interactions is a subfield of the broader field of predicting protein-ligand interactions (PLIs). Generally, there are multiple ways to predict PLIs: (i) Qualitatively, i.e., if a pair of protein and ligand is binding or not, (ii) quantitatively predicting how strong an interaction is measured as binding affinity, and (iii) positionally, where a docking algorithm determines the pose of the ligand relative to the protein [3, 4, 5]. In the latter case, it is typically assumed that the ligand binds to the protein, as most docking algorithms always return a pose, while the first two types of predictors can also correctly predict non-binders. In this work, we focus on qualitative prediction methods for interactions between enzymes and small molecules.

Predicting enzyme–small molecule interactions is challenging due to their inherent complexity caused by the high specificity of enzymes, dynamic and transient interactions with ligands, and conformational flexibility during catalysis [6, 7, 8]. Additionally, enzymes are often multifunctional and promiscuous binders [9, 10]. The evolutionary diversity of enzymes and the lack of high-quality data exacerbate these challenges, making the design of efficient computational methods to predict enzyme functionality even more challenging [7, 8].

Training a model to predict all interaction types (e.g., substrates, regulators, cofactors, and non-interactors) remains challenging due to the lack of comprehensive, curated enzyme–small molecule datasets; consequently, many studies reduce the task to a binary classification problem, distinguishing known, interacting enzyme–small molecule pairs from either experimentally validated [11] or synthetically generated [13, 14, 15, 16, 17, 18] negative samples, where interacting small molecules are primarily understood as substrates. Computational methods for predicting enzyme-substrate interactions thrived recently, evidenced by the publication of several new models and datasets [11, 13, 14, 15, 16].

Since experimental data on non-interacting small molecules is scarce, synthetically generated enzyme–non-interacting pairs are often used to train computational models. Kroll et al. introduced a new dataset categorizing enzyme-small molecule pairs as non-interacting or substrates [13]. Together with the dataset (hereafter called the *ESP dataset*), a deep learning-based model, Enzyme-Substrate Prediction (ESP), has been presented to solve the posed classification task. This dataset has been used in various follow-up studies, including ProSmith and FusionESP [14, 15], which reported remarkable classification performance with AUC values (area under the ROC curve) ranging from 0.956 to 0.972.

*Information leakage* is a phenomenon that can overinflate models’ performance when not properly taken into account. Information leakage (also called *data leakage*) refers to the situation in which a model, during development, has access to information about the evaluation data that is not present in the inference data [12]. In this work, we investigate the influence of information leakage on the performance of three state-of-the-art models: ESP [13], ProSmith [14], and FusionESP [15]. All three models acknowledge the importance of information leakage and report performance drops if evaluated on molecules or drugs unseen in training. They consider two OOD settings: (i) small molecules in the test set that are scarcely or not at all represented in the training set, and (ii) enzymes with 40 *−* 60% and *<* 40% sequence identity to enzymes in the training set. These are typical examples of so-called *inter-sample similarity-induced information leakage*.

In our previous work [19], we investigated in depth the problem of splitting protein-small molecule interaction datasets to minimize information leakage and proposed a total of six OOD settings, of which the above-mentioned OOD tests correspond to only two. Hence, we set out to investigate the three models by re-splitting the original ESP dataset using our recently published DataSAIL [19] and demonstrate that performance declines significantly as inter-sample similarity-induced information leakage decreases further. We show that all three models still rely on inter-sample similarities, namely the presence of similar (in terms of their features) data points in the training and test datasets. If we minimize these similarities, constructing true out-of-distribution splits, models’ performance drops to near-random levels.

## 2 Methods

In the ESP data processing pipeline, experimentally validated interacting pairs of enzymes and small molecules are first clustered by enzyme similarity; the clusters are then randomly split into training and test sets. Importantly, no such clustering is conducted on the side of the small-molecule ligands. However, in ProSmith and Fusion-ESP, the ESP dataset is initially divided into training, validation, and test sets. To ensure consistency, we recomputed this split to train all models. In the following, we refer to this method as the *ESP split* [13]. In addition to that split, we compute several splits generated by DataSAIL (Figure 1): *protein-based* and *ligand-based one-dimensional splits* that split all interactions based on protein or ligand identifiers (I1_L_ and I1_P_, respectively) or similarity clusters (S1_L_ and S1_P_, respectively), and a *two-dimensional split*, S2, that combines S1_L_ and S1_P_ splits, therefore minimizing protein and ligand similarities between splits simultaneously. Because the S2 split removes data, we computed an ESP_S2_ split following the ESP split algorithm, using only the data in the S2 split. This is done to compare the ESP split with the S2 split, the most strict DataSAIL split. In each splitting scheme, we split the data into 70% for training, 10% for validation and hyperparameter tuning, and 20% for testing (Table 3).

**Figure 1.**
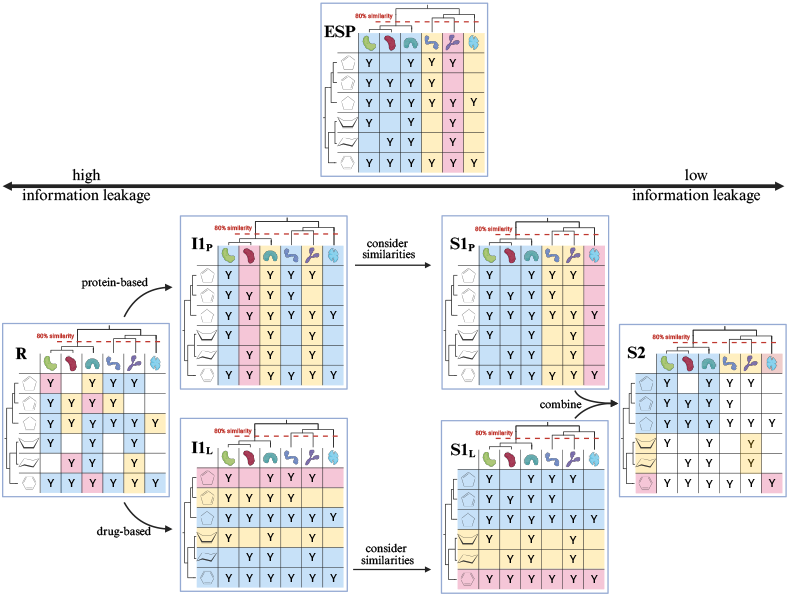
Visualization of data splitting techniques used in this study. The two-dimensional matrix represents the dataset of experimentally verified enzyme-substrate pairs. The symbol “Y” indicates the presence of a measurement, while the dendrograms next to the rows and columns indicate similarities between samples. The dashed red line marks the level beyond which connected nodes have a similarity of 80% or more. Training samples are highlighted in blue, and test samples are in red. Unassignable tiles are left white. In the ESP split, the test set comprises proteins with pairwise similarity to the training data below 80%. We compare this with the six splits below, where we split the dataset either randomly (R), by individual ligand (I1_L_), by individual protein (I1_P_), or by their clusters (S1_L_ and S1_P_). Those splits make sure that no two interactions from different splits share the same (or similar, in the case of S1 splits) ligands (or proteins). In these splits, the data is split only along one dimension (proteins or ligands); the S2 split combines the leakage prevention measures from S1_L_ and S1_P_ at the cost of not assigning all interactions, as some interactions will consist of two molecules in different splits [19].

Theoretically, DataSAIL’s cluster-based splits, S1_L_, S1_P_, and S2, are the most difficult to learn on, because similarity-based splits remove information leakage more rigorously. This has also been previously demonstrated by us for several drug-target interaction datasets[19], but not for the ESP dataset. One major difference between the ESP split and DataSAIL’s S1_P_ split is that the ESP split only ensures that no two proteins in different splits have a similarity above 0.8, while DataSAIL minimizes the similarities between proteins in different splits. Therefore, DataSAIL considers all sequence similarity values, whereas ESP distinguishes values only by whether they are above or below 80% sequence identity [19].

Information leakage can be quantified following [19] for each pair of splits *s* and *s*′, e.g., training and test, of a dataset 𝒟:

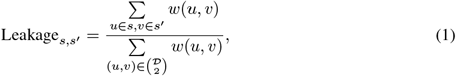

where *u* and *v* are data points in 𝒟, which can be enzymes, substrates, or interactions thereof, and *w*: 𝒟 *×*𝒟 → ℝ is a similarity function between data points in 𝒟. The numerator measures the similarity between distinct splits, while the denominator normalizes by total similarity in the dataset.

Lower leakage values indicate greater dissimilarity between the corresponding splits, resulting in a larger distributional shift between them.

An overview of the models’ architectures and details of the training process can be found in the Supplementary Information.

## 3 Results

We retrained the three models – ESP [13], ProSmith [14], and FusionESP [15] – on the splits described above and performed hyperparameter tuning on the validation sets. All performance metrics calculated here are extracted from the same test sets for all models. The ESP-column of Table 1 shows that our study reproduced the results from the original papers within a minor error margin.

**Table 1:**
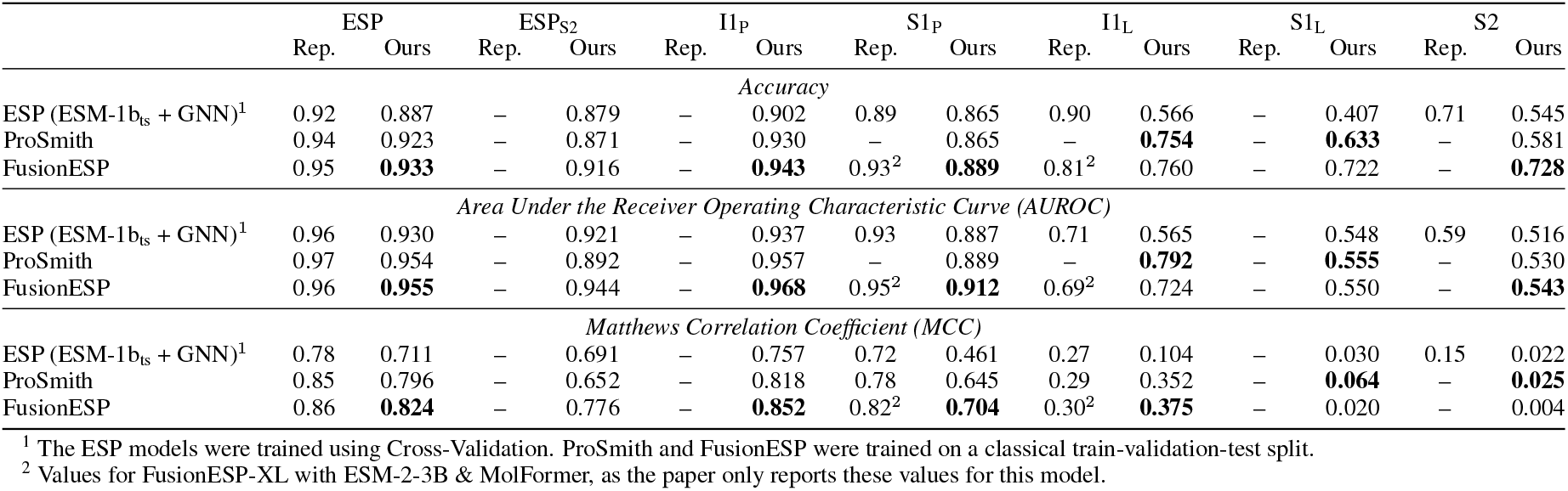
Comparison of performance metrics reported in the original publications and our re-trained models. We computed the Accuracies, the AUROCs, and MCCs on the test set after re-training the models (Ours) and compared them to the values reported (Rep.) in [13, 14, 15]. The metrics in the columns for reported similarity-based cold-protein split (S1_P_-Rep.) refer to the values from the original publications computed on the set of proteins with ≤40% sequence identity to the training set. Similarly, the reported similarity-based two-dimensional cold split (S2-Rep.) is the performance on the subset of interactions in the test set with unseen small molecules and enzyme sequence identity levels of 0 *−* 40% to the training set. The best performance per metric and split is highlighted in bold.

To investigate data leakage in the ESP dataset and assess the models’ generalizability to new enzymes and ligands, we applied various splitting methods using DataSAIL and calculated the leakages using formula 1. First, we compared molecule-similarity leakage (MSL) and protein-similarity leakage (PSL) for different protein-based and ligand-based splits as information leakage calculated using similarity matrices on substrates and enzymes, respectively (Table 2). We observed that splitting a protein-ligand dataset based solely on the protein or the ligands reduces leakage in that axis, but the leakage along the other axis remains high. This can also be observed from the total similarity leakage (TSL), which is computed as MSL + PSL. Overall, ligand-based splits were more effective at minimizing total data leakage, with the S1_L_ split achieving the lowest leakage among the one-dimensional split methods. The lowest overall data leakage was observed in the S2 split, which splits data based on data point similarity along both axes. The reason why protein-based splits consistently show higher data leakage than molecule-based can be the imbalance in the ESP dataset, which includes 12,008 unique proteins and only 1,354 unique small molecules, creating inherent challenges for splitting the data. The enzyme-ligand interaction datasets can be formally represented as a bipartite graph where enzymes and substrates form two disjoint node sets, with edges representing observed interactions. When applying protein-based splitting to this structure, the high connectivity of substrates across enzyme partners frequently creates bridges between training and test partitions, as many substrates interact with multiple enzymes. Their inclusion in the training and test sets across different enzyme partners introduces unavoidable leakage. This architectural constraint explains why protein-based splits consistently show higher data leakage than molecule-based splits in our experiments.

**Table 2:**
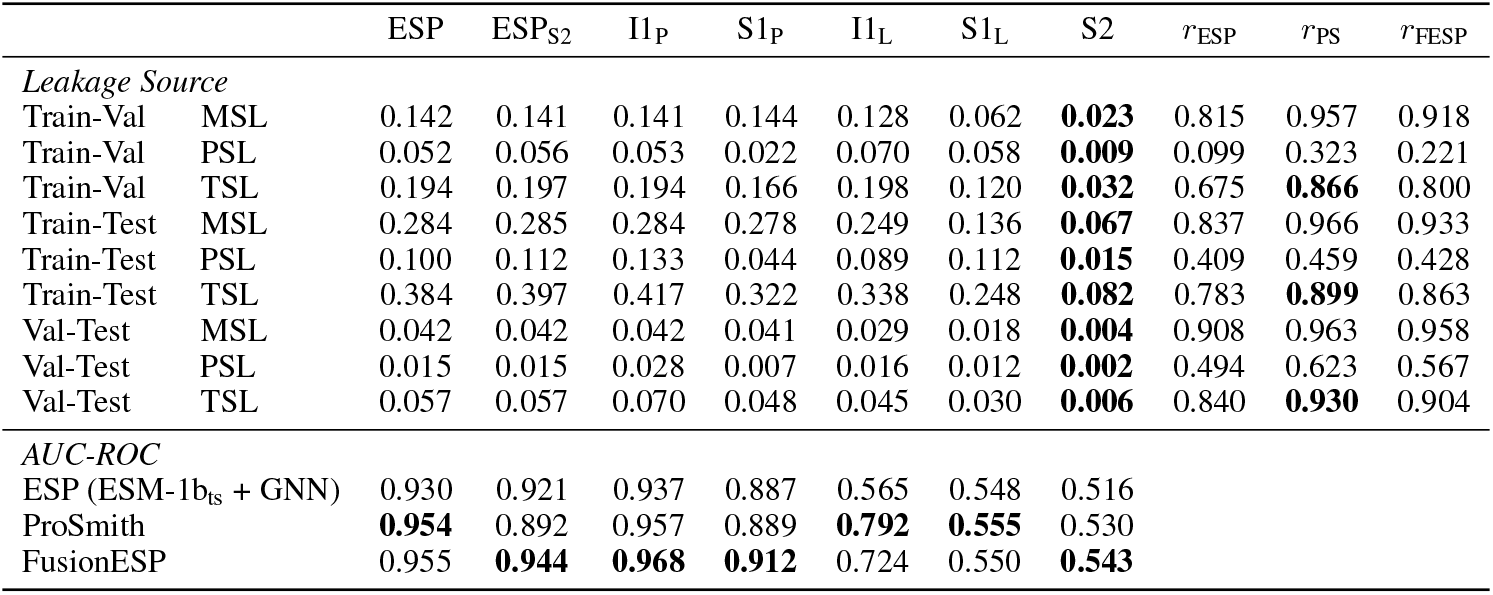
Data leakage and model performances. Comparison of leakages between different subsets, the AUROC of the tested models, and the Pearson correlation coefficient *r* between AUC-ROC and the leakage of the data splits. In bold, we highlighted the lowest leakage values per row, the best model performance per split, and the highest correlation between AUROC and total similarity leakage (TSL).

**Table 3:**
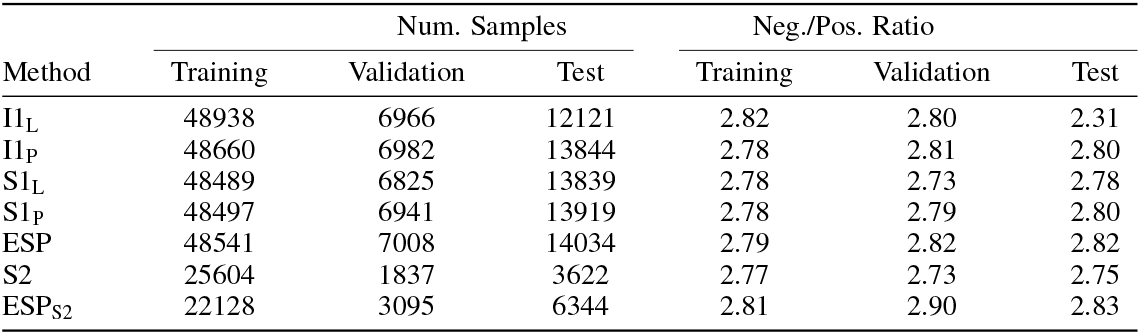
Overview of different split strategies with corresponding sample ratios and negative-to-positive (Neg./Pos.) data ratios.

In the development of the models on the ESP split, information leakage was controlled only along the protein axis, ensuring that no protein in the validation or test sets had more than 80% sequence identity to any protein in the training set. In addition to evaluating the models on the full ESP split test set, the authors also evaluated them on subsets thereof. Two of these subsets are all proteins with 0-40% sequence identity to any sequence in the training set (S1_P_-Rep.), and another one contains only small molecules that were not present in the training set (I1_L_-Rep.). It is important to note that, in the original publications, the model was not retrained for these evaluation settings, whereas we trained a separate model for each splitting technique.

This trend is also reflected in the decreasing performance as splits leak less information and thus become more difficult to memorize (see Table 2, the *r*-columns on the right). Previous studies showed that FusionESP performed best among the three models, with ProSmith as the second best model[15]. Interestingly, in our experiments for the I1_L_ split, ProSmith performs best, and for the S1_L_ split and the S2 split, all models perform almost as a random guess (Table 1).

We observe a similar pattern across the reported Matthews correlation coefficients (MCCs) from the three publications, recalculated MCCs from re-trained models, and MCCs on leakage-reduced splits (Table 1): a decrease of MCC correlates with reduced information leakage. Here, we loosely clustered the reported OOD splits with the most similar DataSAIL splits (e.g., “unseen small molecules”[13] for ESP with S1_L_ and “small molecules not represented in the training set and an enzyme sequence identity level of 0-40% to the training data”[13] with S2). The differences in the MCC values for these OOD splits can be explained by the procedural difference in the algorithms used to create the OOD sets and the DataSAIL splits. While in the original publications, the authors extracted the OOD subsets from the ESP-split test set, we compute splits for each OOD setting individually. This increases the size of the OOD test set but also results in different training and validation sets each time. Additionally, the reported equivalent to our S1_P_ split contains all enzymes with less than 40% sequence identity to any enzyme in the training set. DataSAIL does not provide a fixed upper bound on sequence identity, but the S1_P_ split minimizes the total similarity between the training, validation, and test enzyme sequences.

## 4 Discussion

In this study, we highlight crucial deficiencies in the accepted procedures for training models for enzyme-small molecule interaction prediction that were underexplored before. Existing models are very good at predicting interactions between molecules they encountered during training but struggle to generalize to unseen or OOD data. While generalization to unseen enzymes, even significantly different ones, is good–e.g., FusionESP achieves an AUROC of 0.912 and an MCC of 0.704 on a S1_P_ split–generalization to new small molecules remains poor. This also translates to settings where both the enzyme and the small molecule are new at evaluation time (i.e., S2 split).

Our analysis also fills gaps in previous evaluations (Table 1). It has not previously been investigated how the models perform when the small molecules are structurally dissimilar rather than merely different SMILES strings. Previously, any non-identical molecules have been considered OOD. Here, we extend this to a more fine-grained notion of molecular similarity and show that models cannot generalize significantly beyond near-identical molecules.

These findings also align well with other reports on data leakage in deep learning-based methods for protein-protein interaction prediction [20, 21]. The group showed across multiple works that state-of-the-art deep learning models for protein-protein interaction prediction do not achieve accuracy above 0.65 when information leakage is rigorously removed. Table 1 shows similar results for enzyme substrate prediction models. For the S1_L_ and S2 splits, the deep learning models achieve accuracies (far) below naive baselines. A model always predicting “not interacting” would achieve an accuracy of *∼*0.735 on all splits due to the class imbalance (Table 3).

## Data and Code availability

All data splits we calculated in this study is available on zenodo [28]. The code is available on [GitHub](https://github.com/kalininalab/ESP_Leak_Analysis) and backed up on zenodo [29].

## Author contributions statement

O.V.K conceived the study. V.A.E. and R.J. analysed the data and conducted experiments. R.J. and O.V.K. jointly supervised the study. All authors wrote and reviewed the manuscript.

## Acknowledgments

The authors thank Dr. Alexander Gress for valuable feedback and discussions.

## Supplementary note: Overview of the analyzed models

### Enzyme Substrate Prediction model (ESP)[13]

The ESP model follows the classical architecture of protein-ligand interaction prediction models, with independent encoders for the enzyme and small-molecule components, whose embeddings are concatenated and fed into a prediction head. The enzyme encoder is a modified ESM-1b model[22] in which the authors added an additional 1280-dimensional token representing the entire enzyme, learning a task-specific representation of the enzyme. Small molecules are either represented as 1024-dimensional ECFP descriptors with a radius of 3 or using a graph neural network. The prediction head is a gradient-boosting model (Figure 2a). The authors report very good performance in testing (AUC: 0.956), but observed a significant decline in performance when evaluating only the out-of-distribution parts of the test set (AUC: 0.71; MCC: 0.27; Table 1).

**Figure 2.**
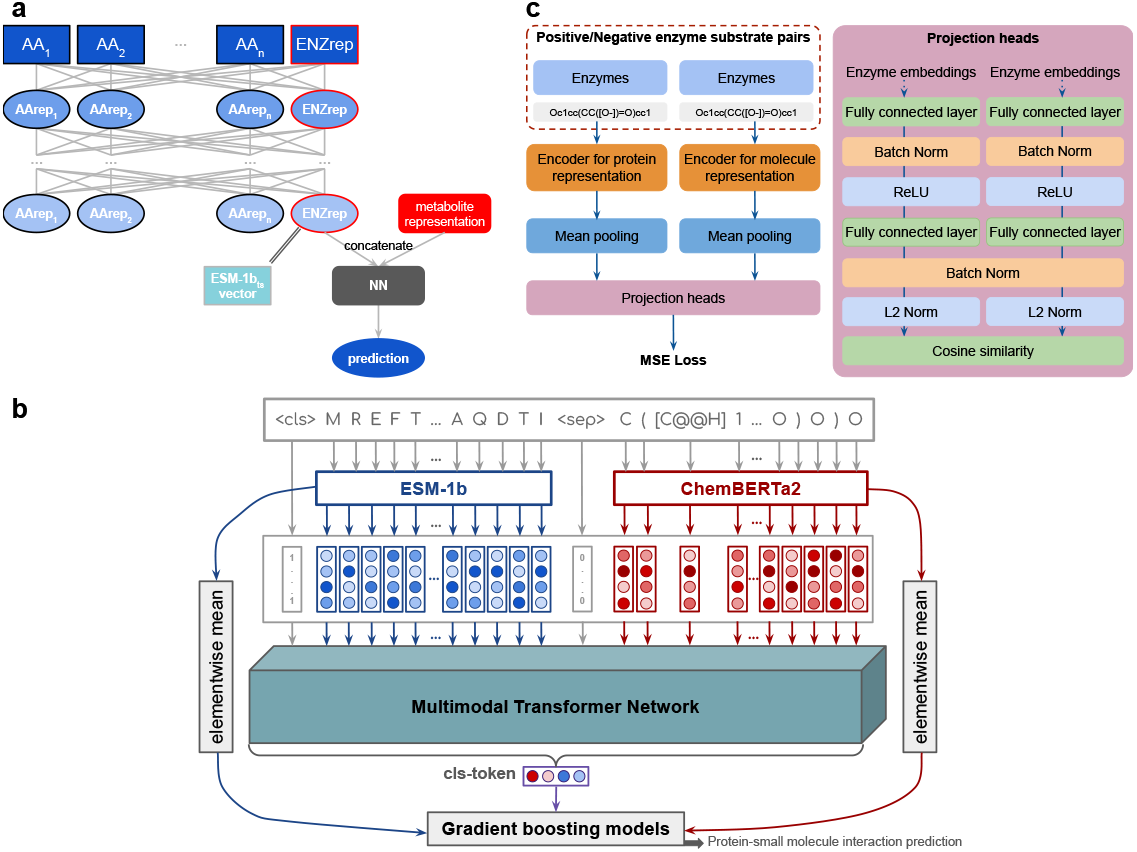
Schemata of tested models. Here, we depict the workflows of the different models investigated. All graphics are inspired by the depictions in the original publications and closely follow their structure. **a** shows ESP, **b** shows ProSmith, and **c** shows FusionESP.

### PROtein-Small Molecule InTeraction, Holistic model (ProSmith)[14]

This model processes token-level ESM-1b protein embeddings and ChemBERTa2[23] ligand embeddings in a multimodal transformer. Both embeddings are concatenated with a 0-vector as a separator in between and prepended with a unit-vector as a cls-token. From this transformer, the cls-token embedding is extracted and paired with the mean-pooled embeddings from ESM-1b and ChemBERTa2. The CLS token alone, the combined ESM-1b and ChemBERTa2 embeddings, or their combination with the CLS token are then used as input to a gradient-boosting prediction head. In this study, we focus on the model trained using the combination of all three representations. (Figure 2b). ProSmith is pre-trained on IC50 values from the BindingDB dataset [24], and then task-specifically fine-tuned on the downstream datasets Davis[25] and ESP. To ensure that the downstream validation and test splits are truly OOD, all proteins and ligands from those splits were removed from the pre-training data in the BindingDB. Like ESP, ProSmith was separately evaluated on the OOD parts of the test set and showed clear improvement over ESP models to an MCC of 0.29 (Table 1).

### FusionESP[15]

FusionESP also follows the general idea of having two independent encoders for proteins and molecules. It updates the previously used ESM-1b protein encoder to an ESM2 encoder[26] and substitutes ChemBERTa2 with MoLFormer[27]. In this study, we focus on the FusionESM model with the ESM2-650M model over the variant utilizing the ESM2-3B model. On the forward pass, the mean-pooled embeddings are separately passed through a projection head comprising two streams, each with two rounds of a dense layer, batch normalization, and either ReLU activation or L2 regularization. The prediction is then calculated as the cosine similarity between the two resulting 128-dimensional embeddings (Figure 2c). This similarity is then optimized using a mean-squared error loss function. This has the effect that the embeddings of proteins and ligands lie in the same latent space, and that the embeddings of binding molecules are close to each other, while those of non-binding pairs are pushed apart. Furthermore, embeddings of proteins with similar binding patterns are projected close together. The evaluation shows that FusionESP both outperforms ESP and ProSmith, with the gap to ProSmith smaller than to ESP. For OOD subsets of the test data, FusionESP’s margin over ProSmith further increases to AUC 0.8185 (see Table 1).

All models were retrained using Python 3.12 and PyTorch 2.8 on a single workstation with a NVIDIA RTX 3090 GPU with 24GB RAM and 16 CPUs with 128GB RAM. The data were downloaded from the ESP GitHub repository and the linked Zenodo archive [13]. For ProSmith and FusionESP, we also followed the instructions from their respective GitHub repositories [14, 15].

## References

[1] Jeremy M. Berg, John L. Tymoczko, Gregory J. Gatto, Lubert Styrer. Biochemistry Springer 524–560, 2012.

[2] Inger Andersson. Catalysis and regulation in Rubisco. Journal of experimental botany 59(7):1555–1568 (2008).

[3] Oleg Trott and Arthur J. Olson. AutoDock Vina: improving the speed and accuracy of docking with a new scoring function, efficient optimization, and multithreading. Journal of computational chemistry 31(2):455–461 (2010).

[4] Antoine Lacour, Hamza Ibrahim, Andrea Volkamer, and Anna KH Hirsch. DockM8: An All-in-One Open-Source Platform for Consensus Virtual Screening in Drug Design. preprint DOI: 10.26434/chemrxiv-2024-17k (2024).

[5] Gabriele Corso, Hannes Stärk, Bowen Jing, Regina Barzilay, and Tommi Jaakkola. “Diffdock: Diffusion steps, twists, and turns for molecular docking. preprint DOI: 10.48550/arXiv.2210.01776 (2022).

[6] Robert Callender and R. Brian Dyer. The dynamical nature of enzymatic catalysis. Accounts of chemical research 48(2):407–413 (2015).

[7] Gordon G. Hammes, Stephen J. Benkovic, and Sharon Hammes-Schiffer. Flexibility, diversity, and cooperativity: pillars of enzyme catalysis. Biochemistry 50(48):10422–10430 (2011).

[8] Ioannis G. Riziotis, António J. M. Ribeiro, Neera Borkakoti, and Janet M. Thornton. Conformational variation in enzyme catalysis: a structural study on catalytic residues. Journal of Molecular Biology 434(7):167517 (2022).

[9] Olga Khersonsky and Dan S Tawfik. Enzyme promiscuity: a mechanistic and evolutionary perspective. Annual review of biochemistry 79(1):471–505 (2010).

[10] Yin-Ming Kuo, Ryan A. Henry, and Andrew J. Andrews. Measuring specificity in multisubstrate/product systems as a tool to investigate selectivity in vivo. Biochimica et Biophysica Acta (BBA)-Proteins and Proteomics 1864(1):70–76 (2016).

[11] Dante A. Pertusi, Matthew E. Moura, James G. Jeffryes, Siddhant Prabhu, Bradley Walters Biggs, and Keith E. J. Tyo. Predicting novel substrates for enzymes with minimal experimental effort with active learning. Metabolic engineering 44:171–181 (2017).

[12] Judith Bernett, David B. Blumenthal, Dominik G. Grimm, Florian Haselbeck, Roman Joeres, Olga V. Kalinina, and Markus List. Guiding questions to avoid data leakage in biological machine learning applications. Nature Methods 21(8):1444–1453 (2024).

[13] Alexander Kroll, Sahasra Ranjan, Martin KM Engqvist, and Martin J. Lercher. A general model to predict small molecule substrates of enzymes based on machine and deep learning. Nature communications 14(1):2787 (2023).

[14] Alexander Kroll, Sahasra Ranjan, and Martin J. Lercher. A multimodal Transformer Network for protein-small molecule interactions enhances predictions of kinase inhibition and enzyme-substrate relationships. PLOS Computational Biology 20(5):e1012100 (2024).

[15] Zhenjiao Du, Weimin Fu, Xiaolong Guo, Doina Caragea, and Yonghui Li. FusionESP: Improved Enzyme–Substrate Pair Prediction by Fusing Protein and Chemical Knowledge. Journal of Chemical Information and Modeling 65(6):2806–2817 (2025).

[16] Zhiwei Nie, Hongyu Zhang, Hao Jiang, Yutian Liu, Xiansong Huang, Fan Xu, Yonghong Tian, Jie Chen, and Wen-Bin Zhang. Multi-purpose enzyme-substrate interaction prediction with progressive conditional deep learning. prerpint DOI: 10.21203/rs.3.rs-5516445/v1 (2024).

[17] Wenjia Qian, Xiaorui Wang, Yuansheng Huang, Yu Kang, Peichen Pan, Chang-Yu Hsieh, and Tingjun Hou. Deep Learning-Driven Insights into Enzyme–Substrate Interaction Discovery. Journal of Chemical Information and Modeling 65(1):187–200 (2024).

[18] Haiyang Cui, Yufeng Su, Tanner J. Dean, Tianhao Yu, Zhengyi Zhang, Jian Peng, Diwakar Shukla, and Huimin Zhao. Enzyme specificity prediction using cross attention graph neural networks. Nature (2025): 1–3.

[19] Roman Joeres, David B. Blumenthal, and Olga V. Kalinina. Data splitting to avoid information leakage with DataSAIL. Nature Communications 16(1):3337 (2025).

[20] Judith Bernett, David B. Blumenthal, and Markus List. “Cracking the black box of deep sequence-based protein–protein interaction prediction.” Briefings in Bioinformatics 25, no. 2 (2024): bbae076.

[21] Timo Reim, Anne Hartebrodt, David B. Blumenthal, Judith Bernett, and Markus List. Deep learning models for unbiased sequence-based PPI prediction plateau at an accuracy of 0.65. Bioinformatics 41(Supplement_1):i590–i598 (2025).

[22] Alexander Rives, Joshua Meier, Tom Sercu, Siddhart Goyal, Zeming Lin, Jason Liu, Demin Guo, Myle Ott, C. Lawrence Zitnick and Rob Fergus. Biological structure and function emerge from scaling unsupervised learning to 250 million protein sequences. Proceedings of the national academy of sciences 118(15), e2016239118 (2021).

[23] Walid Ahmad, Elana Simon, Seyone Chithrananda, Gabriel Grand and Bharath Ramsundar. Chemberta-2: Towards chemical foundation models. preprint DOI: 10.48550/arXiv.2209.01712 (2022).

[24] Tiqing Liu, Yuhmei Lin, Xin Wen, Robert N. Jorissen and Michael K. Gilson BindingDB: a web-accessible database of experimentally determined protein–ligand binding affinities. Nucleic acids research 35(suppl_1) D198–D201 (2007).

[25] Mindy I. Davis, Jeremy P. Hunt, Sanna Herrgard, Pietro Ciceri, Lisa M. Wodicka, Gabriel Pallares, Michael Hocker, Daniel K. Treiber, and Patrick P. Zarrinkar. Comprehensive analysis of kinase inhibitor selectivity. Nature biotechnology 29(11) 1046–1051 (2011).

[26] Zeming Lin, Halil Akin, Roshan Rao, Brian Hie, Zhongkai Zhu, Wenting Lu, Nikita Smetanin, Robert Verkuil, Ori Kabeli, Yaniv Shmueli, Allan Dos Santos Costa, Maryam Fazel-Zarandi, Tom Sercu, Salvatore Candido, and Alexander Rives Evolutionary-scale prediction of atomic-level protein structure with a language model. Science 379(6637) 1123–1130 (2023).

[27] Fang Wu, Dragomir Radev and Stan Z. Li Molformer: Motif-based transformer on 3d heterogeneous molecular graphs. Proceedings of the AAAI Conference on Artificial Intelligence 37(4) 5312–5320 (2023).

[28] Information Leakage in Enzyme Substrate Prediction. Zenodo DOI: 10.5281/zenodo.18786394.

[29] Information Leakage in Enzyme Substrate Prediction. Zenodo DOI: 10.5281/zenodo.18788610.

